# State-dependent geometric transformations in the mouse hippocampus support fear generalization without loss of discriminability

**DOI:** 10.64898/2026.05.01.722221

**Authors:** Hung-Tu Chen, Yosif Zaki, Denise J. Cai, Matthijs A. A. van der Meer

## Abstract

Learning from aversive experiences often generalizes beyond the context in which they occurred. In rodents, a strong aversive event can induce retrospective memory linking (RLI), whereby fear generalizes to a previously neutral context encountered days earlier. Although prior work has shown that RLI is associated with increased co-activity of hippocampal CA1 neurons across neutral and aversive contexts, it remains unclear how broader representational changes support generalization without affecting the ability to discriminate between contexts. Here, we reanalyzed calcium imaging data from dorsal CA1 during RLI to examine how hippocampal representational geometry changes during fear generalization. Using robust, non-parametric measures of population similarity, we show that in mice exhibiting RLI, the representation of the neutral context not only changes over time but becomes more similar to the aversive context during recall. Beyond this similarity increase, we provide evidence for a higher-dimensional geometric transformation consistent with a shared “fear” operation that can be applied across contexts while preserving their identity. Crucially, these two representational signatures dissociate by behavioral state: similarity to the aversive context emerges during freezing, whereas a shared transformation is expressed during active exploration. Together, these findings demonstrate that hippocampal representations support retrospective fear generalization through state-dependent geometric transformations, highlighting representational geometry as a key computational mechanism to resolve the apparent tension between generalization and discriminability.

## Introduction

How do aversive experiences become linked with past memories to influence subsequent behavior? A notable feature of aversive experiences is their tendency to generalize to other situations or contexts beyond the one in which the experience originally occurred (Bouton, 2004; Maren et al. 2013; Dunsmoor & Paz, 2015). For instance, the phenomenon of *retrospective memory linking*, (RLI) begins when an animal encounters an aversive event in a given context, and subsequently shows fear in the context in which the event occurred (“aversive context”). Interestingly, if the shock intensity is sufficiently high, the animal will perceive a previously neutral environment (i.e. where no shock occurred) as similarly threatening when re-encountered later -- retrospectively linking the aversive event with the neutral environment encountered days earlier (Cai et al. 2016, Zaki et al. 2025; Figure 1).

**Figure 1:**
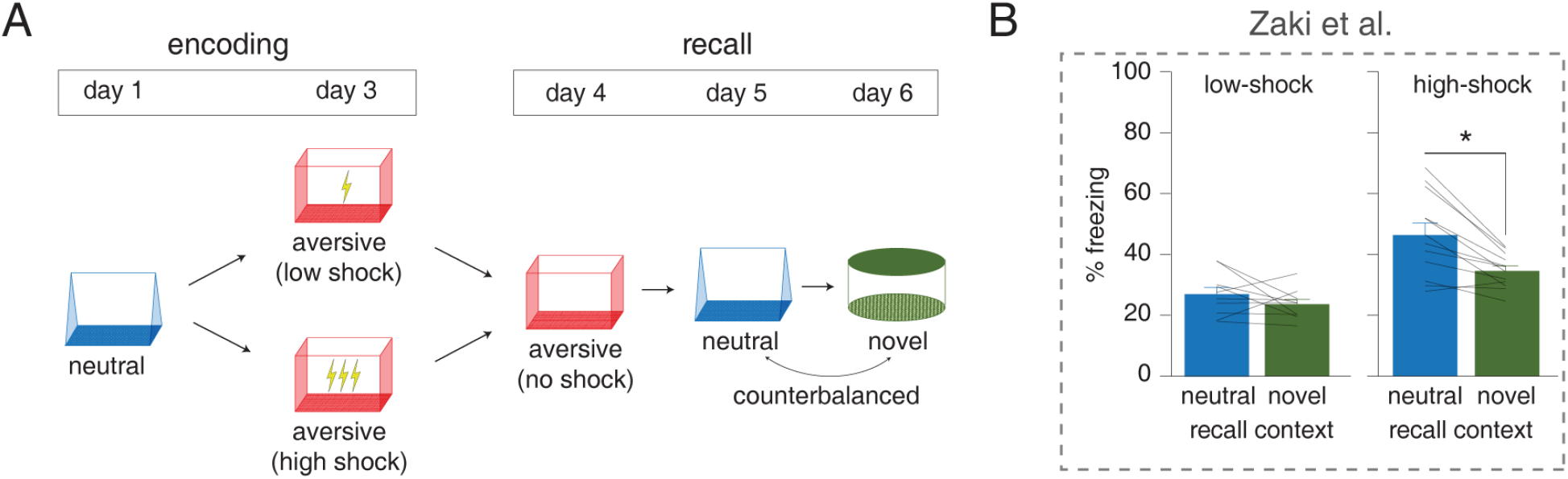
Retrospective memory linking: strong aversive experience induces retrospective memory-linking to a previously neutral context, as shown in Zaki et al. (2025) **(A)** Schematic of the retrospective memory-linking experiment. On day 1, mice were introduced to a neutral context. After two days, separate groups of mice received either 3 low-intensity shocks (0.25mA) or 3 high-intensity shocks (1.5mA) in an aversive context. On the following day, both groups were tested in an aversive context. Over the next two days, testing occurred in both the neutral and a novel context in a counterbalanced order. **(B)** Comparison of freezing levels during the recall sessions in the neutral and novel contexts for both groups. Mice exposed to the high-shock condition exhibited significantly increased freezing behavior in the neutral context compared to the novel context (p < 0.01 for Wilcoxon signed-rank test), while mice in the low-shock group did not show this difference (p = 0.25). This result highlights the key behavioral phenomenon of fear memory being linked retrospectively to a different, previously neutral context. Errorbars indicate SEM across subjects. Adapted from Zaki et al. 2025.

Contextual fear and its generalization, such as occurs in RLI, depends on the hippocampus (Kim & Fanselow 1992; Anagnostaras et al. 2001). What neural representations and computations could underlie RLI? There are two main ideas: the *first idea* is to keep distinct neural representations for the neutral (N) and aversive (A) contexts intact (conventionally, with orthogonal activity patterns, i.e. global remapping; Figure 2a), but update a downstream association between the neutral content and shock (Figure 2b). This approach leaves the ability to discriminate between N and A intact (i.e. no confusion about what environment you are in); however, without some representational similarity between A and N it is unclear by what logic or mechanism N would be targeted for association with shock after experience in A (which is represented by an independent neural ensemble). The second idea is to update the neural representation of N to be more like A, i.e. generalizing by representational similarity (Figure 2c). This idea provides a clear computational and mechanistic basis for generalization: representational overlap is induced by associative learning (e.g. TCM, Howard & Kahana 2002 and other associative learning theories; McClelland & Rumelhart 1985, O’Reilly & Rudy 2001) and the extent of representational overlap determines generalization. However, in this account, a drawback is that representational overlap comes at the cost of reduced discriminability between N and A (i.e. becoming confused about which environment you’re in).

**Figure 2:**
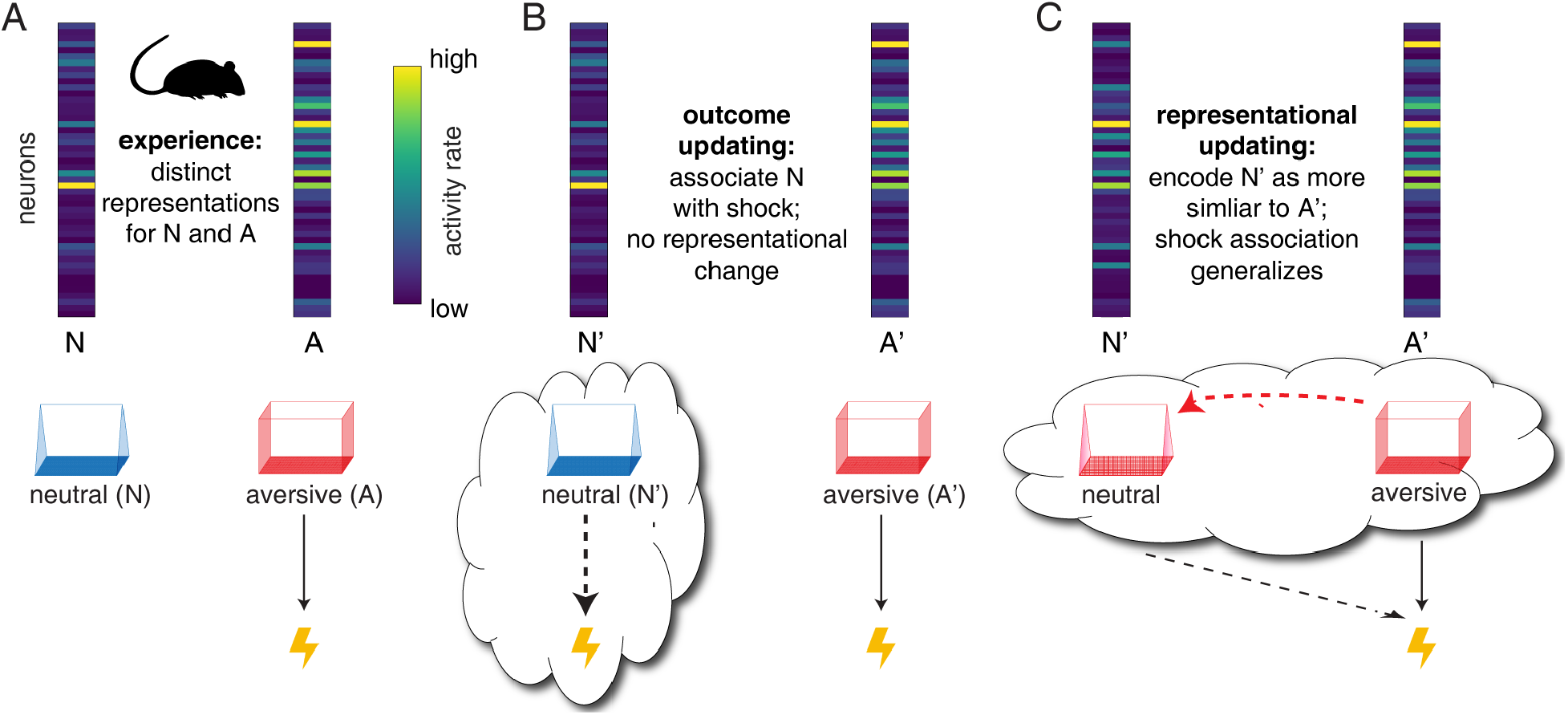
Framework for the neural representations and computations that may underlie retrospective memory linking. **(A)** When an animal experiences a sufficiently aversive event (e.g. footshock), it learns to associate fear not only with the context where the event occurred (the “aversive context”, A) but also with a previously experienced neutral context, N. This retrospective memory linking (RLI) phenomenon presents a puzzle because initially, hippocampal activity patterns representing A and N are distinct (top row). **(B)** Outcome updating hypothesis: in this scenario, the animal maintains stable, distinct representations of the aversive and neutral contexts, while associating the neutral context with the aversive event in a downstream brain structure (e.g. amygdala) without changing the core representation of the neutral context. Thought bubbles highlight the association being updated in this hypothesis. **(C)** Representational updating hypothesis: Alternatively, the hippocampal representation of the neutral context may be dynamically updated to resemble the aversive context, facilitating memory linking by making the neural representations of the two contexts more similar. The neutral context becomes associated with the aversive event by virtue of its similarity (representational overlap) with the aversive context.

Thus, the “downstream-association” and “representational-overlap” ideas highlight a tension: on the one hand, without some kind of representational similarity between A and N, it is unclear by what logic or on what mechanistic basis generalization can happen. On the other, generalization by representational similarity seemingly carries a cost of decreased discriminability. Prior work has shown that retrospective memory linking is associated with an increase in the number of neurons that are active in both N and A (e.g. Cai et al. 2016; Rashid et al. 2016; Zaki et al. 2025); however, if such co-activity were the sole cause of generalization, then it also induces forgetting (i.e. ability to discriminate between N and A). Importantly, an extensive literature across tasks and species has shown that adaptive generalization does not equal forgetting (Blough 1967; Ghirlanda & Enquist 2003). How can this tension between generalization and discriminability be resolved?

One plausible solution we investigate here is that the answer lies in **representational geometry**: when contextual representations are conceived as multi-dimensional patterns of activity, rather than as one-dimensional “overlap”, it is possible to independently encode the identity of a context (N vs. A) and whether or not it is associated with shock (Figure 3; see Fusi et al. 2016 for a review of related “mixed selectivity” ideas). To test how the representational geometry of hippocampal activity changes during RLI, we re-analyzed the calcium imaging dataset from Zaki et al. (2025) where mice generalize fear from a recent aversive experience to a previously neutral experience. We first identified hippocampal CA1 neurons active across all recording sessions (i.e. neutral and aversive encoding and recall sessions) and examined the geometry of their activity patterns during memory linking. Extending previous results that found an increase in neurons co-active in both N and A, we first show that the neural representation of the neutral context becomes more similar to the aversive context in terms of representational similarity. Interestingly, by assessing the geometry of the representational space, we found that the direction of these shifts is influenced by the animal’s behavioral state: during freezing, the neutral context representation becomes more similar to the aversive context, whereas during active exploration, it aligns with a “common fear” operation, transitioning from a fear-free to a fearful context without a change in ensemble similarity. These findings offer the first direct evidence of how hippocampal representational geometry (i.e. not just “co-activity” or overlap in active neurons) shifts when fear responses acquired in an aversive context generalize to a past neutral context.

**Figure 3:**
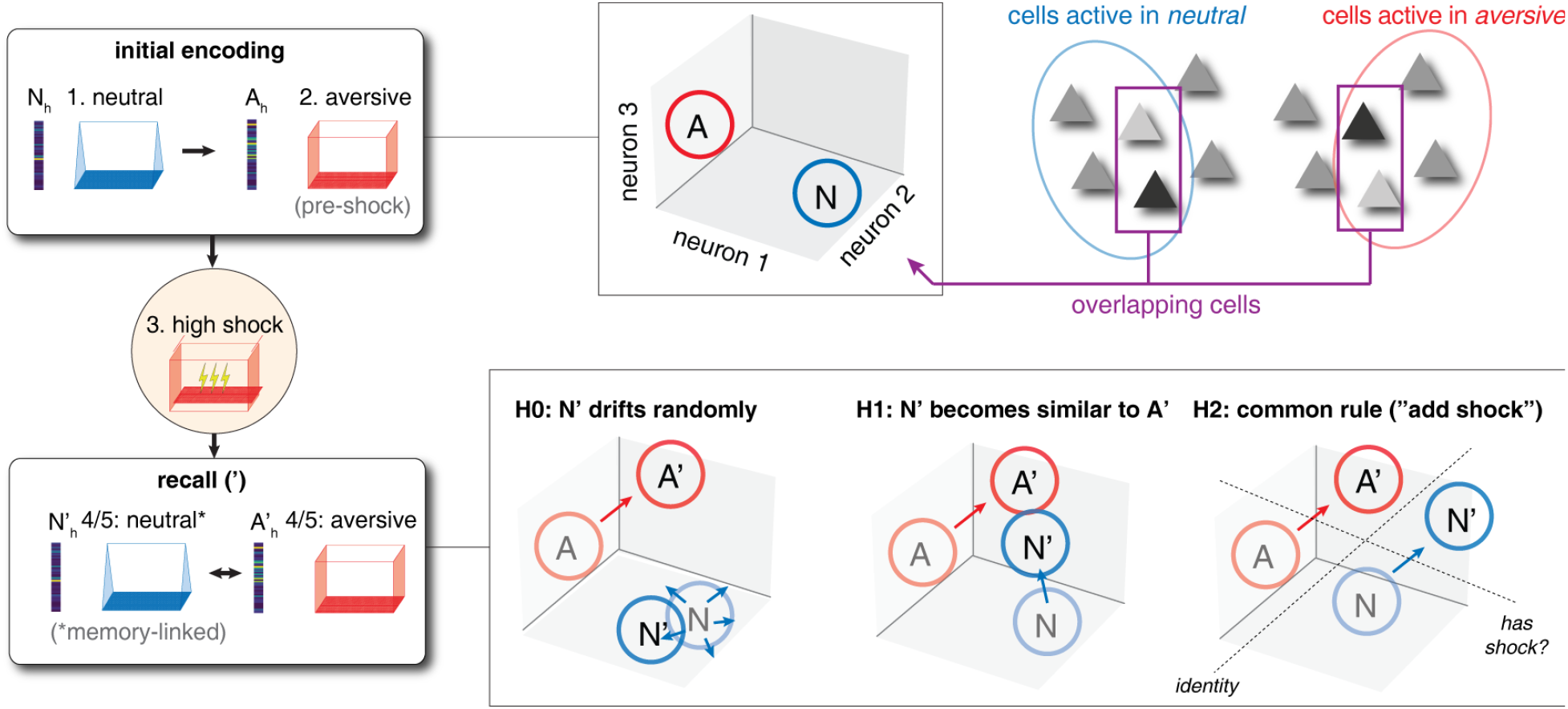
Hypothesized representational changes in neural activity space underlying memory linking. **Top row:** During initial encoding of experience, separate hippocampal activity patterns for the neutral (N, step 1) and aversive (A, step 2) contexts are formed. Previous work has examined overlap between these patterns (i.e. the proportion of cells active in both environments, binary yes/no participation); in this study, we examine changes in the *pattern* of activity in cells that are active in both contexts. **Bottom row**: Following high-intensity shock in the aversive environment (step 3, highlighted circle) mice show freezing in the previously neutral environment (steps 4 and 5; order of exposure to N and A is counterbalanced across mice). We consider two hypotheses for changes in neural activity patterns that may underlie this generalization. The null hypothesis is that the neural representation for N drifts randomly (H0, left). The “similarity increase” hypothesis (H1, middle) is that the neutral environment representation becomes more similar to the aversive representation (note the N to N’ change brings N’ closer to A’ than N is to A). The second “common rule” hypothesis (H2, right) holds that N and A change in similar ways, such that the ability to decode identity is preserved, but a downstream decoder could distinguish both A vs A’ and N vs N’ with the same decision boundary (“do I expect shock in this environment?”) orthogonal to the identity-coding dimension.

## Results

To understand how hippocampal representations are modified during retrospective memory-linking, we re-analyzed the calcium imaging dataset from Zaki et al. (2025). In this study, mice with calcium imaging recordings in the dorsal CA1 region first experienced a neutral context (N). Two days later, separate groups of mice were introduced to an aversive context (A) paired with either a low or high-intensity shock (see Figure 1 for a schematic of the timeline). Both groups were subsequently tested in the aversive, neutral and novel contexts. Notably, the high-shock group (but not the low-shock group) exhibited increased freezing during recall in the neutral context compared to the novel context, indicating that fear memory had been retrospectively linked to the previously neutral context.

Previous accounts of the retrospective memory-linking phenomenon have shown that the aversive memory becomes retrospectively linked with a neutral context through the offline reactivation of hippocampal CA1 neurons that are co-active between the aversive and neutral memories (Zaki et al., 2025). However, this account leaves a major question unresolved: What specific representational changes beyond co-activity (a simple, one-dimensional measure) accompany memory linking? Specifically, for those neurons that are active in both contexts, do the activity *patterns* in the aversive and neutral contexts (i.e. population vectors) stay stable or change randomly (Figure 3; null hypothesis “H0”, consistent with the idea that retrospective memory linking occurs through downstream associations, Figure 2b)? Alternatively, do the aversive and neutral activity patterns become more similar to each other (Figure 3; hypothesis “H1”; Figure 2c)? Or, does the neutral activity pattern change in such a way that it maintains the same representational distance to the aversive experience, but can be decoded as aversive due to a parallel geometry or common rule (Figure 3; hypothesis “H2”, resolving the confound between discriminability and generalization that exists for a one-dimensional measure of similarity like the number of co-active neurons)?

To investigate these possibilities, we compared the pattern similarity between population vectors consisting of neurons active across both environments (i.e. vectors of length `n_neurons`, containing the mean activity of each neuron, one for each environment and experimental condition to be compared; see *Methods* for details) between encoding and recall sessions for both neutral and aversive contexts (neutral: N and N’; aversive: A and A’; see Figures 2 and 3 for schematics). To ensure that population activity from these different recording sessions was discriminable to begin with, we first trained a linear discriminant analysis (LDA) model to assess separability between sessions (Table 1). Our results confirmed that all pairs of neutral and aversive sessions were distinguishable from chance levels (range: 68-88% accuracy; Wilcoxon signed rank test, p < 0.05 compared to 50%). Importantly, decoding accuracy between the neutral and aversive encoding sessions (*N vs. A*) was similar across low-shock and high-shock groups (p = 0.22 for Wilcoxon rank-sum test), as expected; here and for subsequent analyses, we only used pre-shock data for the encoding sessions to avoid potential confounds due to behavioral differences associated with the shock.

**Table 1:**
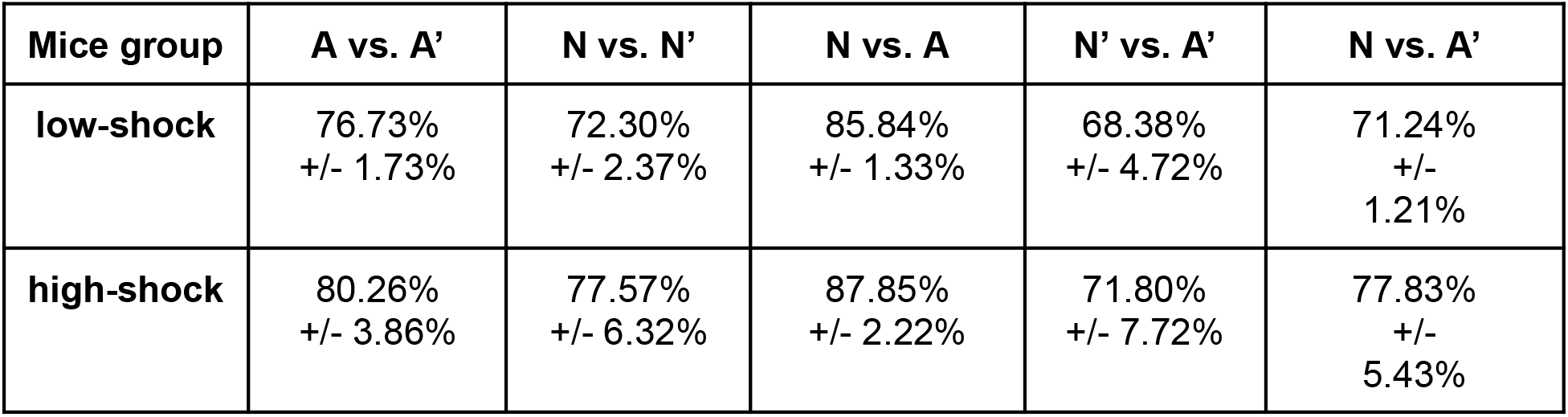
Decoding accuracy using linear discriminant analysis (LDA) for neutral and aversive sessions in low-shock and high-shock mice. We evaluated decoding accuracy between each pair neutral and aversive sessions across both low-shock and high-shock groups of mice. **A** and **N** indicate the aversive and the neutral *encoding* sessions respectively (only pre-shock data was used to minimize behavioral confounds). **A’** and **N’** indicate the aversive and the neutral *recall* sessions. The LDA model was trained on 80% of the data, with the remaining 20% used for testing, and decoding accuracy was averaged over five distinct train/test splits. All pairs of neutral and aversive sessions were separable from chance level (Wilcoxon signed rank test, p < 0.05 compared to 50%).

| Mice group | A vs. A' | N vs. N' | N vs. A | N' vs. A' | N vs. A' |
| --- | --- | --- | --- | --- | --- |
| low-shock | 76.73%<br>+/- 1.73% | 72.30%<br>+/- 2.37% | 85.84%<br>+/- 1.33% | 68.38%<br>+/- 4.72% | 71.24%<br>+/- 1.21% |
| high-shock | 80.26%<br>+/- 3.86% | 77.57%<br>+/- 6.32% | 87.85%<br>+/- 2.22% | 71.80%<br>+/- 7.72% | 77.83%<br>+/- 5.43% |

Next, we sought to determine the most reliable representational similarity measure for these data by comparing several similarity metrics including non-parametric Kendall’s tau correlation, and parametric measures such as cosine similarity, Pearson correlation, and Euclidean distance (Figure S1). As a testbed, we focused on the representational similarity between the neutral encoding session and the pre-shock period of the aversive encoding session; ideally, across different sessions and subjects, this representational similarity should not depend on incidental factors (e.g. the number of recorded neurons) and should be robust to skewed activity distributions that can arise from including variable numbers of high-firing interneurons. Comparing various different measures, our analyses showed that Kendall’s tau correlation was the most robust measure, as it showed minimal correlations of representational similarity with both neuron count (r = 0.06) and the proportion of high-firing neurons (r = 0.04). This finding aligns with previous studies showing that parametric similarity measures are more susceptible to biases introduced by skewed neural firing rate distributions or voxel activations (Nili et al., 2014; Walther et al., 2016). Thus, Kendall’s tau was chosen for the representational similarity tests that follow.

### How does the representation of the neutral context change from encoding to recall?

To begin our representational similarity analysis, we first extended the findings of Zaki et al. using Kendall’s tau on the ensemble activity of neurons co-active across environments. In the low-shock group, both the neutral and aversive contexts showed a similar degree of change from encoding to recall (p = 0.44, Wilcoxon signed-rank test; left column, Figure 4a), even though the encoding and recall sessions for the neutral context were further apart in time compared to the aversive context. However, in the high-shock group, the correlation between encoding and recall was significantly lower for the neutral context compared to the aversive context (p < 0.05, right column, Figure 4a; analogous to Zaki et al’s result). This lower correlation between encoding and recall in the high-shock group suggests that in mice that display retrospective memory-linking, the neutral context undergoes more substantial changes from encoding to recall than the aversive context (and more changes than in the low-shock group that does not generalize fear).

**Figure 4:**
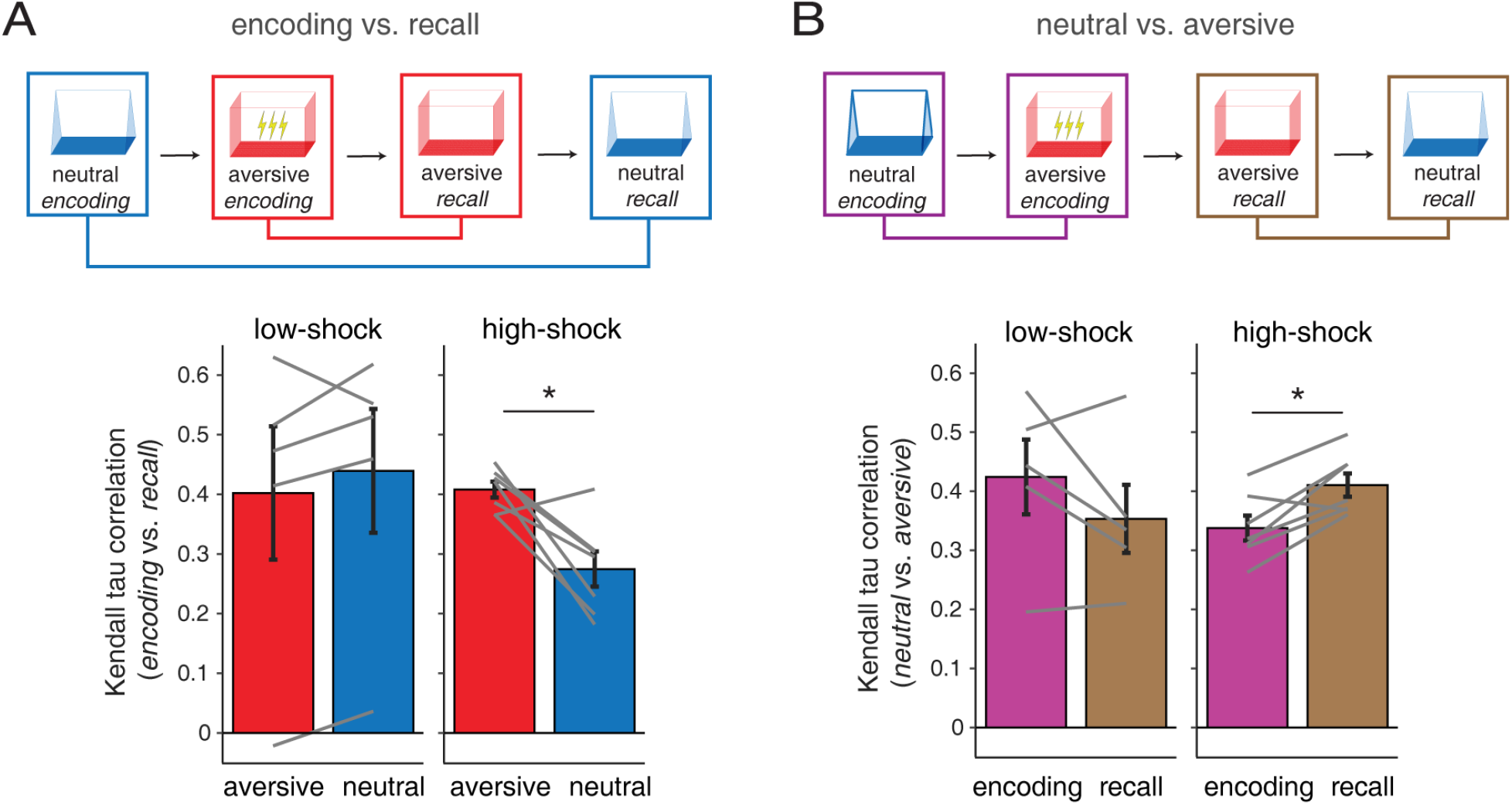
High shock causes the neutral context to change and become more similar to the aversive context compared to low shock. **(A)** We compared population activity patterns between the encoding and recall sessions of the same context (i.e. neutral encoding vs. neutral recall, blue; and aversive encoding vs. aversive recall, red). In the high-shock group, population activity patterns in the neutral context were significantly less correlated between encoding and recall compared to the aversive context (p < 0.05 for Wilcoxon signed-rank test), indicating larger representational change for the neutral compared to aversive context. In contrast, the low-shock group showed no significant difference between neutral and aversive encoding-to-recall correlations (p = 0.44). **(B)** Next, we compared population activity between the neutral and aversive contexts during either encoding or recall sessions. In the high-shock group, population activity vectors for the neutral and aversive contexts became more correlated during recall compared to encoding (p < 0.05), indicating increased representational similarity following high shock. Conversely, the low-shock group showed no significant change in neutral-to-aversive correlations between encoding and recall sessions (p = 0.31). Error bars indicate SEM across subjects.

One way in which changes in the neutral context representation could support memory-linking between the two contexts is that the population activity patterns in the neutral and aversive contexts becomes more similar in the high-shock group (*Hypothesis 1* in Figure 3). Consistent with this prediction, we observed a significant increase in neutral-to-aversive correlations from encoding to recall in the high-shock group (p < 0.05, Wilcoxon signed-rank test; right column, Figure 4b). In contrast, no significant change in the similarity between these two context representations was found in the low-shock group (p = 0.31, left column, Figure 4b). These results show that for the high shock group, the representation of the neutral context not only changes from encoding to recall, but also becomes more similar to the aversive context, supporting Hypothesis 1.

The above analysis shows that in retrospective memory linking, the increase in neural similarity between the aversive and neutral environments extends beyond an increase in the number of co-active neurons to also include increased pattern similarity in those co-active cells. However, several issues remain unresolved: (1) as discussed in the *Introduction*, simple one-dimensional measures like co-activity and pattern similarity confound generalizability with discriminability, and (2) based on the extensive literature on “place cells” the moment-to-moment neural representation of space and context is known to depend on behavioral state (e.g. immobility vs. active exploration; Colgin, 2016; McNaughton et al., 1983; O’Keefe & Dostrovsky, 1971; Wilson & McNaughton, 1994), raising the question to what extent the above pattern similarity results are state-specific. To address these issues, we perform a representational similarity analysis to distinguish between different behavioral states and different representational geometries

### In which direction does the neutral context representation change?

Specifically, we investigated which *direction* the neutral context activity shifts in to support memory-linking (see Figure 3 for a schematic): does the previously neutral context become linked to (1) the aversive context during recall (*context shift*), or become modified with (2) a common “fear” operator (*common rule*)? To distinguish between these possibilities, we compared observed changes in the neutral context representation with different hypothesized directions, while separately analyzing freezing and non-freezing periods to account for different behavioral states. To do this, we computed Kendall’s tau correlations between the population-activity difference vector from the neutral encoding session to the neutral recall session, and each of the following: 1) the difference vector from neutral encoding to aversive recall (*context shift*), and 2) the difference vector from pre-shock periods during aversive encoding to aversive recall (*common rule*; see Figure 3 for schematic). We then compared these correlations across different behavioral states (freezing vs. active exploration) between the low-shock and high-shock groups. A significant difference between groups would suggest that the neutral context undergoes representational changes in the direction of the hypothesized shift. In addition, for both groups, we compared these correlations to a control distribution obtained by shuffling context labels to assess whether the observed correlations were due to random neural activity patterns.

Our results showed significant differences between the high-shock and low-shock groups during freezing periods. Specifically, in the high-shock group, the neutral context shifted significantly more toward the representation of the aversive event than in the low-shock group (i.e. context shift; p < 0.05, Wilcoxon rank-sum test with contrast, Figure 5a). In contrast, during active exploration, a significant shift was only observed in the direction of a common “fear” transformation of the neutral context (p < 0.05; Figure 5b). Furthermore, across both behavioral states (context shift and common rule), we found significant differences between freezing and active exploration states in the high-shock group (p < 0.05, Wilcoxon signed-rank test with contrast). Specifically, the context shift effect was only observed during freezing, whereas the common rule effect was only observed during active exploration. Importantly, these shifts were significantly greater than those observed in the shuffled control distributions, ruling out the possibility that the effects were due to random fluctuations in hippocampal activity (all p < 0.05, bootstrap test). This double dissociation provides evidence for behavioral state-dependent representational geometry changes -- an increase in representational similarity between aversive and neutral environments during freezing, and a fear transformation that preserves representational distance between environments during active exploration.

**Figure 5:**
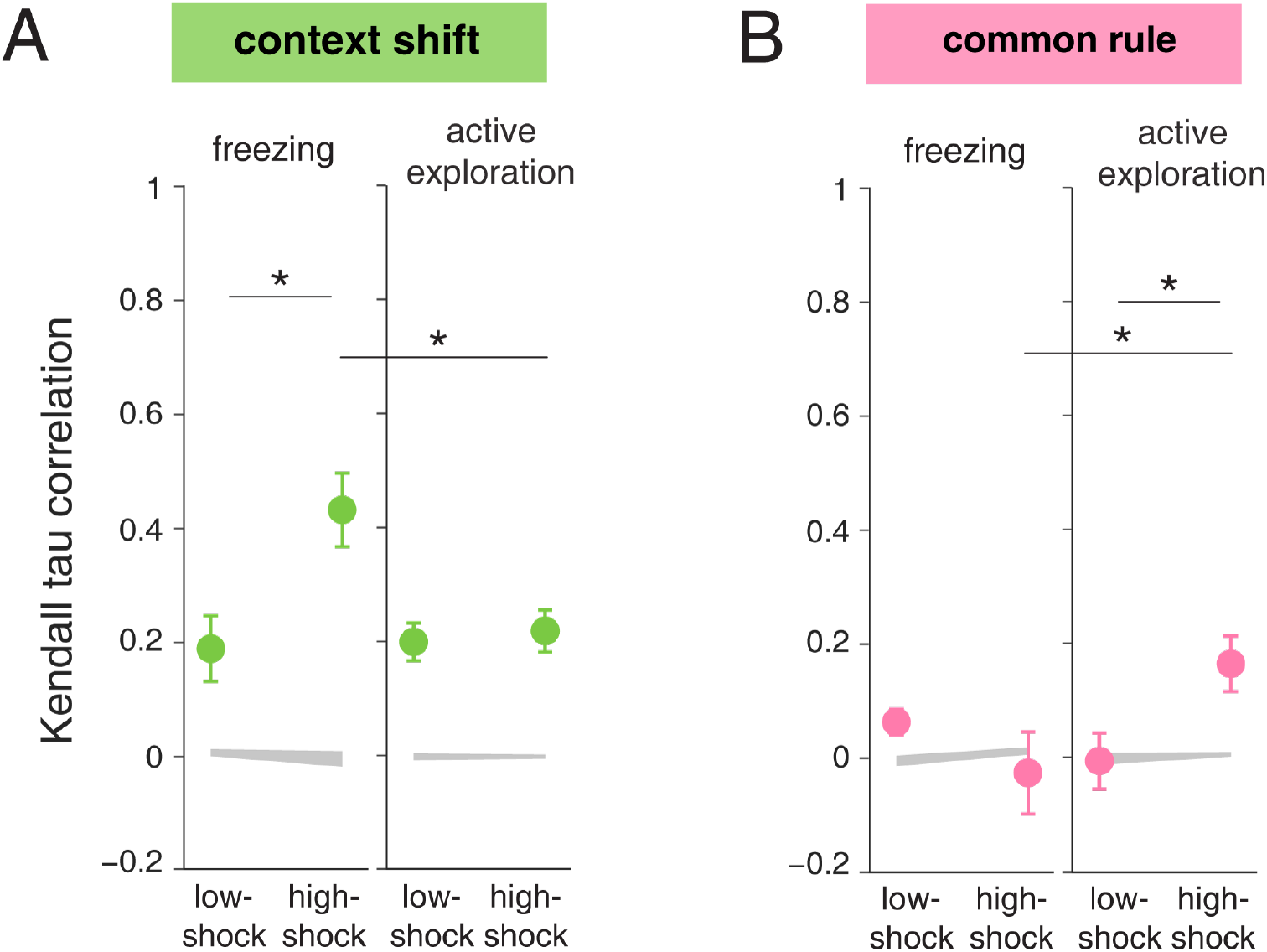
Memory linking involves distinct mechanisms of representational change depending on behavioral state. **(A) Context shift:** We computed correlations between the population-activity difference vector from the neutral encoding session to the neutral recall session and the difference vector from neutral encoding to *aversive recall*. In the high-shock group, there was a significantly stronger shift in the neutral context representation toward the aversive experience during freezing states (p < 0.05, Wilcoxon rank-sum test with contrast), but no significant shift was observed during active exploration (p = 0.94). Furthermore, a significant difference between freezing and active exploration was observed only in the high-shock group (p < 0.05, Wilcoxon signed-rank test with contrast), but not in the low-shock group (p = 0.72). (**B) Common rule**: We correlated the population-activity difference vector (encoding vs. recall) in the neutral context with the difference vector from *pre-shock periods during aversive encoding to aversive recall*. During active exploration, memory-linked mice exhibited a significant “common fear” transformation in neutral context representations (p < 0.05). Again, only the high-shock group showed a significant difference between freezing and active exploration (p < 0.05). Error bars indicate SEM across subjects. Shaded regions represent the correlations in a shuffled control distribution (see *Methods* for details).

## Discussion

In this study, we first extend previous work showing that the hippocampal (CA1) representation of a neutral context becomes more similar to an aversive context, specifically in the group of mice that generalize fear to that neutral context (retrospective memory linking or RLI; Cai et al. 2016, Zaki et al. 2025). This similarity increase occurs not just in terms of an increase in the number of neurons co-active across the neutral and aversive contexts, as shown previously (Zaki et al. 2025) but as we now show, in terms of neural pattern similarity in co-active neurons. This result bolsters the idea that aversive experience causes dynamic updates to the hippocampal representation of context, and bridges the distinct literatures memory allocation and engrams using calcium imaging (which typically examine co-activity or overlap) on the one hand, and place cell ensemble recordings using electrophysiology (which typically examine population vectors or patterns, as we do here) on the other.

However, one-dimensional measures of neural similarity (whether co-activity or population vector correlation) are unlikely to provide a full explanation of generalization behavior such as occurs in RLI. This is so because if the brain were to rely on such 1-D representations only, then generalization would be confounded with discriminability: when two contexts are represented more similarly so that freezing behavior generalizes, the ability to discriminate between those contexts for other behaviors necessarily decreases proportionally. In reality, the level of flexibility shown in typical generalization behaviors suggests that the brain does not confound these things (Blough 1967; Ghirlanda & Enquist 2003). Thus, we also tested the geometrical possibility that a higher-dimensional “mixed” pattern encodes context identity along one axis and the association with shock along an orthogonal axis (Figure 3, Hypothesis 2).

We found support for this common-rule idea, showing that the pattern transformation (i.e. a vector in neural activity space) that occurs between encoding and recall of the aversive environment can be applied to the initially neutral environment to predict how this environment will change with behavioral generalization of fear to this environment. This idea can be thought of as an “add association with shock” operation that is applied to the aversive environment because of experience, and to the neutral environment because of generalization (i.e. a common rule that can be applied to different environments). While the statistical evidence for this common-rule idea was robust, we noted that the magnitude of the effect was relatively modest (Kendall’s tau correlations of ∼0.15 compared to chance level of 0); we discuss some possible reasons for this below.

A striking feature of these two complementary representational changes associated with generalization, i.e. the increase in pattern similarity between neutral and aversive contexts, and the existence of a common-rule operation to reflect the shock association, was their double dissociation depending on behavioral state. Pattern similarity between the two contexts was only apparent when mice were immobile, whereas common-rule coding was only visible during movement. This is an interesting observation in its own right, reminiscent of known differences in hippocampal place and context coding depending on behavioral and neural state (e.g. theta rhythms associated with movement and exploration vs. irregular activity during wakeful rest vs. delta rhythms associated with freezing). It is also helpful in ruling out the ever-present possibility that any reported changes in neural representational similarity are actually due to changes in behavioral similarity (e.g. the animals freezing more in the previously neutral context after generalization, making behavior potentially more similar to that in the aversive context).

Further investigation of the significance of this double dissociation would benefit greatly from improved temporal resolution. The behavioral-state-dependent shifts in the neutral context representation likely reflect two distinct cognitive processes. During freezing states, memory retrieval, potentially involving sharp-wave ripples, may dominate, causing the neural representation to shift toward the aversive experience or context (Buzsáki, 2015; Ólafsdóttir et al., 2018; Wu et al., 2017). Conversely, during active exploration, memory encoding processes, possibly linked to theta oscillations, might transform the neutral context from fear-free to fearful (Backus et al., 2016; Buzsáki & Moser, 2013; Skaggs et al., 1996). To validate these hypotheses, higher-resolution electrophysiological recordings are needed to more accurately discriminate between the physiological states and cognitive processes involved. Similarly, the need to average the neural activity population vectors across all locations in each environment likely limited our ability to detect strong signals for changes in neural representations such as the common-rule operation.

A related avenue for further work is to determine whether similarity increases truly reflect a change in context, or instead recall of a different context or even a specific episodic event (e.g. being shocked); this will require fine temporal resolution to distinguish e.g. different theta cycles which are known to “flicker” between different environments and trajectories (Jezek et al., 2011; Kay et al., 2020), and/or sharp wave-ripples which can reflect specific episodes (Buzsaki 2015; Berners-Lee et al. 2022). In any case, the interplay between shifts in representational geometry and behavioral state we observed hints at a richer understanding of the neural basis of generalization across environments.

More generally, our results also highlight the importance of placing CA1 representational geometry within the broader hippocampal circuit. Extensive work has implicated dentate gyrus and CA3 in pattern separation and pattern completion, respectively, suggesting that similar experiences can either be orthogonalized or linked through subregion-specific computations (Leutgeb et al. 2007; McHugh et al. 2007; Neunuebel & Knierim 2014; Kesner & Rolls 2015). CA1, the focus of the present analysis, is therefore unlikely to “solve” the generalization problem alone. Instead, CA1 population geometry may reflect the combined influence of upstream DG/CA3 computations as well as behavioral state. Future experiments recording across hippocampal subfields during retrospective memory linking will be needed to determine whether increased similarity, common-rule transformations, and preserved contextual discriminability emerge locally within CA1 or are inherited from earlier stages of hippocampal processing and inputs.

## Methods

### RESOURCE AVAILABILITY

#### Materials availability

This study did not generate new unique reagents.

#### Data and Code Availability

Calcium imaging data used in this manuscript is available at (https://github.com/denisecailab/RetrospectiveMemoryLinkingData_2024). Code necessary and sufficient for reproducing all results and figures in this manuscript is publicly available on Github (https://github.com/hungtuchen/RLI_colab).

### EXPERIMENTAL MODEL AND SUBJECT DETAILS

This study used experimental data from a previously published dataset (Zaki et al. 2025), which included 14 adult C57BL/6J wild-type mice from Jackson Laboratories, aged 12 to 15 weeks at the beginning of behavioral training. These mice were chronically implanted with Miniscope for calcium imaging recordings. Full subject details are described in Zaki et al. (2025).

### METHOD DETAILS

#### Behavioral task

We conducted new analyses on the Zaki et al. (2025) dataset, comprising calcium imaging recordings of hippocampal CA1 neurons from mice performing a contextual fear conditioning task. Initially, mice were exposed to the *neutral* context for 10 minutes to explore. Two days later, they were placed in the *aversive* context and allowed to explore for 2 minutes. Following this, they received a 2-second foot shock, either 0.25mA (low shock) or 1.5mA (high shock). A second shock of the same duration and intensity was administered one minute after the first, and a third shock followed one minute after the second. Thirty seconds after the third shock, the mice were returned to their home cages. Over the subsequent three days, the mice were tested in the previously encountered *aversive* context first, followed by tests in the *neutral* and completely *novel* contexts, with the order of these latter two tests counterbalanced.

#### Criteria for inclusion of data

Only subjects with at least 10 neurons recorded consistently across sessions (both encoding and recall sessions of neutral and aversive contexts) were included in the analysis, resulting in 13 out of 14 total subjects. Additionally, neurons with activity rates exceeding five standard deviations above the population mean were excluded from the analysis.

#### Data preprocessing

Full recording and preprocessing details are described in Zaki et al. (2025). Briefly:

##### Calcium imaging data

Calcium imaging was recorded at 30 frames per second. We used the open-source pipeline Minian to extract calcium transients from the raw data (Dong et al., 2022). Pre-processing steps included removing background fluorescence and sensor noise, followed by motion correction. Next, putative cell bodies were identified and processed using a constrained non-negative matrix factorization algorithm, which separated the video data into spatial footprints for each cell and a matrix of calcium transients. These transients were then deconvolved to estimate the timing of each calcium event. For cross-session analyses, cells were registered using the open-source toolbox, CellReg, which matches cells across sessions based on the spatial correlations of nearby cell footprints (Sheintuch et al., 2017).

##### Freezing behaviors

Behavioral data were processed using the open-source tracking pipeline, ezTrack, to measure freezing behavior (Pennington et al., 2019). To synchronize these behavioral data with calcium imaging data, the behavior recordings were first aligned to an idealized template based on a perfect 30 Hz sampling rate. The calcium imaging data were then aligned with the behavioral frames by matching each behavior frame to its closest corresponding calcium imaging frame.

### QUANTIFICATION AND STATISTICAL ANALYSIS

All analyses and statistical tests were conducted using Python, utilizing the NumPy, SciPy,, Scikit-Learn, and Pingouin libraries. Error bars represent the standard error of the mean (SEM), and data points with error bars indicate the mean values. Statistical significance was assessed using Wilcoxon signed-rank tests for within-group comparisons and Wilcoxon rank-sum tests for between-group comparisons. Significant effects of interaction were followed with post-hoc testing with the use of contrasts.

#### Representational change analysis

To investigate how neural activity patterns in the neutral context change during memory linking, we used Kendall’s tau correlation to measure the similarity of mean population activity across different session pairs. Only cells that were consistently active across all sessions were included in the analysis. The analysis consisted of three main tests:

#### Encoding vs Recall (Figure 4a)

Following the approach in Zaki et al. (2025), we evaluated whether the neural representation of a specific context (e.g. neutral or aversive) changes from the encoding to recall session. This was done by correlating the mean population activity vectors consisting of co-active cells between these sessions, i.e. neutral encoding-neutral recall, and aversive encoding-aversive recall. Significance tests were performed within each group for both the neutral and aversive contexts, and the results were compared between the low-shock group (which did not exhibit memory linkage) and the high-shock group (which did exhibit memory linkage).

#### Neutral vs Aversive (Figure 4b)

To assess if the representations of the neutral and aversive contexts become more similar from encoding to recall, we correlated the mean population vectors of co-active cells of the neutral context with that of the aversive context, separately for encoding and recall, i.e. neutral encoding-aversive encoding, and neutral recall-aversive recall. Significance tests were conducted on these correlations within each group across encoding and recall sessions, followed by a comparison of the results between the two groups.

#### Representational geometry test (Figure 5)

This test aimed to determine the directional shifts in neural representations during memory linking. We compared the actual change in the neutral context representation (i.e. the difference between the population activity vector of neutral encoding and that of neutral recall) with vectors representing different hypothesized directional changes:

- **Context shift:** the actual change vector was correlated with the vector from the neutral encoding to the aversive recall representation. A significant difference from the baseline would suggest that the neutral context shifts toward the most recent representation of the aversive *context*.
- **Common rule:** the actual change vector was correlated with the vector describing the change from pre-shock periods in aversive encoding to the aversive recall session. A significant difference from the baseline would indicate that the neutral context undergoes a *common fear transformation*, shifting from a fear-free to a fearful state.
- **Random change:** the above hypothesized vectors from the low-shock group served as the baseline. If no significant differences were observed when comparing the high-shock group to this baseline, it would suggest that the representational changes occur randomly and are unrelated to the aversive experience. Additionally, we compared the correlations in the actual data with a shuffled control distribution (obtained through 1000 context-label shuffles) to ensure that observed correlations were not due to random neural activity patterns.

Behavior factors such as immobility are known to impact hippocampal activity and prevent a clear comparison of different representational change vectors. To address this, our analysis included separate evaluations for freezing and non-freezing periods.

#### Session discriminability analysis

To investigate whether neural activity in one session is discriminable from activity in other sessions, for each pair of neutral and aversive sessions we trained a linear discriminant analysis (LDA) model to decode the population activity. Specifically, we trained the LDA model to classify the session to which each time bin belonged, using 80% of the data for training and 20% for testing. Decoding accuracy was averaged across five different train/test splits. We also confirmed the consistency of these results by replicating the analysis with a support vector machine (SVM) model with a linear kernel and a logistic regression model, both of which produced comparable outcomes.

## Acknowledgements

We thank Jeremy Manning, John Murray, Caleb Kemere and David Redish for their helpful feedback and discussion. This work was supported by NIMH R01 MH123466 to MvdM, DP2 MH122399, R01 MH120162, Brain Research Foundation Award, Klingenstein-Simons Fellowship, NARSAD Young Investigator Award, McKnight Memory and Cognitive Disorder Award, One Mind-Otsuka Rising Star Research Award, Hirschl/Weill-Caulier Award, Mount Sinai Distinguished Scholar Award, and Friedman Brain Institute Award, to DJC; NIMH F31MH126543 to YZ.

## Author contributions

HTC, MvdM: conceived research; HTC: wrote analysis code, performed data analysis, generated figures; YZ: collected data; HTC, YZ, DJC, MvdM: interpreted data analysis; HTC & MvdM: wrote manuscript with comments from YZ & DJC.

## Supplementary Information

**Figure S1:**
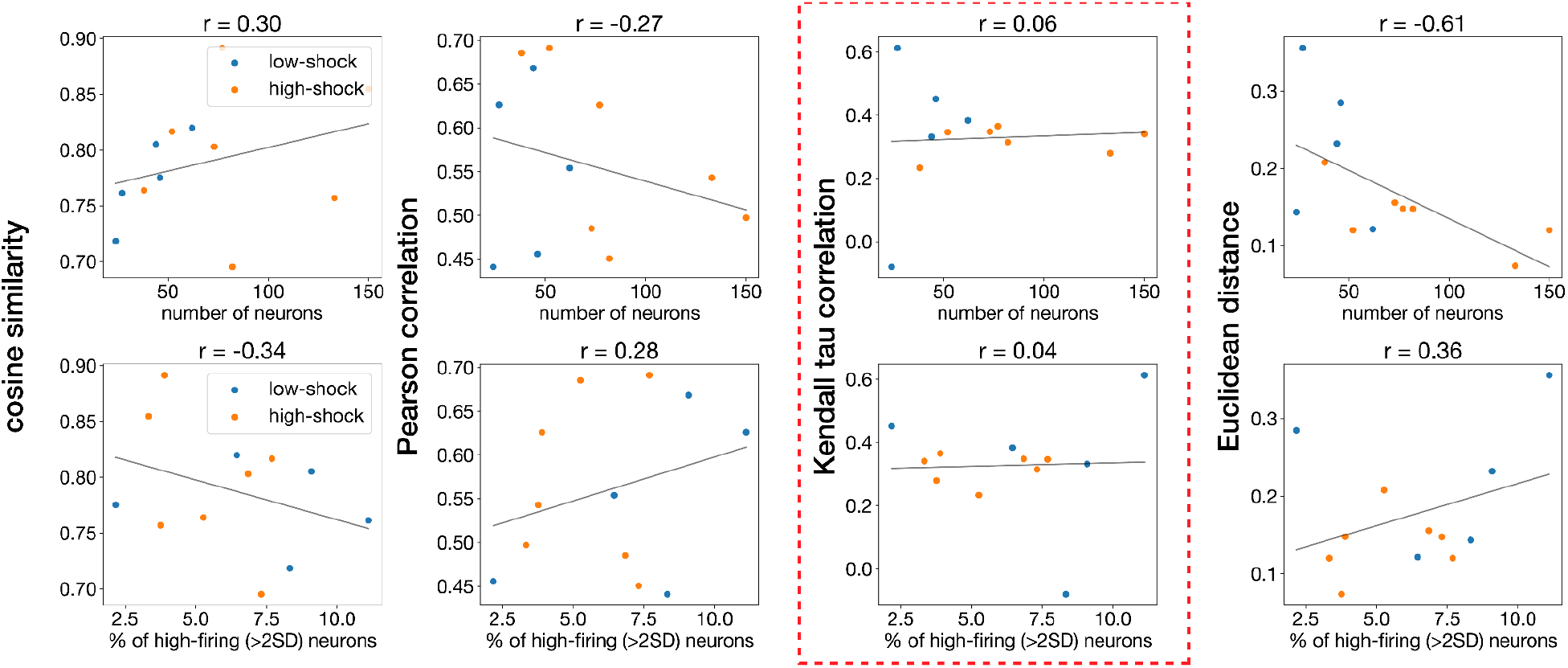
Kendall’s tau correlation outperforms other parametric distance measures. We compared several representational similarity measures, including the non-parametric Kendall’s tau correlation and parametric methods such as cosine similarity, Pearson correlation, and Euclidean distance. We focused on the similarity between neural activity during the neutral encoding session and the pre-shock periods of the aversive encoding session, as these periods are unrelated to shock events and thus provide an unbiased comparison for both the low- and high-shock groups. Each measure was correlated with the number of recorded neurons (top row) and the proportion of high-firing neurons (bottom row), with each point indicating the correlation or distance for an individual subject. Kendall’s tau correlation was the most robust measure, with minimal correlation with both neuron count (r = 0.06) and the proportion of high-firing neurons (r = 0.04).

